# Heart rate changes related to risky selections and outcomes in a rat gambling task

**DOI:** 10.1101/2024.04.21.590481

**Authors:** Fumiya Fukushima, Atsushi Tamura, Nahoko Kuga, Takuya Sasaki

**Affiliations:** Department of Pharmacology, Graduate School of Pharmaceutical Sciences, Tohoku University, 6-3 Aramaki-Aoba, Aoba-Ku, Sendai 980-8578, Japan; Department of Neuropharmacology, Tohoku University School of Medicine, 4-1 Seiryo-machi, Aoba-Ku, Sendai 980-8575, Japan

## Abstract

Risk-taking behavior is crucial for animals to increase their potential outcomes and is considered to alter arousal states in the brain and body represented by heart rates. In this study, we monitored changes in heart rate as rats performed a gambling task in which they selected either a certain outcome with 100% probability (sure option) or a probabilistic double outcome with 50% probability (risky option). We found that when rats selected risky options, they exhibited significantly greater decreases in their instantaneous heart rates immediately before selection than when they selected certain options. In addition, we observed significantly larger increases in instantaneous heart rates when the rats were informed of the larger number of outcomes after selecting the risky options than after selecting the sure options. These results demonstrate that animals can dynamically alter their instantaneous heart rates in response to risky selection and outcomes.

## INTRODUCTION

Risk-taking behavior involves adaptive actions or decisions that include uncertainty about potential benefits, loss, and danger. During risk-taking behaviors, heart rates representing psychophysiological arousal levels have been shown to change when playing various gambling games, including blackjack, (Anderson & Brown, 1984); poker machines, (Leary & Dickerson, 1985); fruit machines, (Griffiths, 1993); and horse racing, (Coventry & Norman, 1997), while change patterns of gambling-related heart rates vary significantly among individuals, related to their gambling frequency or addiction levels ((Leary & Dickerson, 1985; Coulombe *et al*., 1992; Griffiths, 1993; Lole *et al*., 2014)). At fine temporal scales, instantaneous heart rates have been shown to dynamically decrease during the anticipation of outcomes resulting from gambling (Goudriaan *et al*., 2006; Lole *et al*., 2012) and prominently increase or decrease after wins or losses, respectively (Coventry & Hudson, 2001; Goudriaan *et al*., 2006; Lole *et al*., 2012).

As risk-taking behavior is common across species, from rodents to humans, experimental rodent models have been extensively utilized to evaluate decision-making in risky situations, which contributes to understanding the neural basis underlying risk-taking behavior (Ishii *et al*., 2012; Orsini *et al*., 2015; Barrus & Winstanley, 2016; Tremblay *et al*., 2017). However, whether and how animals change their heart rates in response to risky situations remains unknown. To address this question, we monitored the heart rates of rats engaged in a gambling task with risky selections and outcomes.

## Materials and Methods

### Ethical approvals

All experiments were approved by the Committee on Animal Experiments of Tohoku University (approval number: 2023, PhA-007), and were performed in accordance with the NIH guidelines for the care and use of animals.

### Animals

Male 10-to-15-week-old Long Evans rats (SLC, Shizuoka, Japan) with a preoperative weight of 300–400 g were housed under conditions of controlled temperature and humidity (22 ± 1ºC, 55 ± 5%) in a vivarium, maintained on a 12:12-h light/dark cycle (lights off from 8 pm to 8 am) with access to food and water provided ad libitum. Once behavioral training began, food and water were restricted. All rats were housed individually.

### Behavioral setup

The experiments were conducted in a sound-attenuated box (30 × 40 × 30 cm) equipped with a fan for ventilation (Fig. 1A). All behavioral experiments were performed at a light intensity of 1 lx. On one wall of the box, a nose-poke hole (hole 1) was located at the center, and a red LED was positioned above hole 1. Two nose-poke holes (left: hole 2; right: hole 3) were located on each side of hole 1, and a green LED was positioned above holes 2 and 3. A nozzle for delivering a water drop into the water port was located on the opposite wall, and a blue LED was positioned above the nozzle. The delivery of the water drops triggered a clicking sound from the nozzle, which served as a notification to the rats that the water had been dispensed. To detect nose poking and water consumption, infrared photoreflectors were attached as sensors at appropriate positions. Each device was connected to a computer via a Digital I/O card, and was controlled using an in-house software program (based on Python). Depending on the sensor inputs, LED illumination and water feeding were automatically regulated and digitized and timestamped using a laptop computer. Because all the apparatuses were controlled by a computer, it was not necessary for the experimenter to handle the rats after the task began.

**Figure 1.**
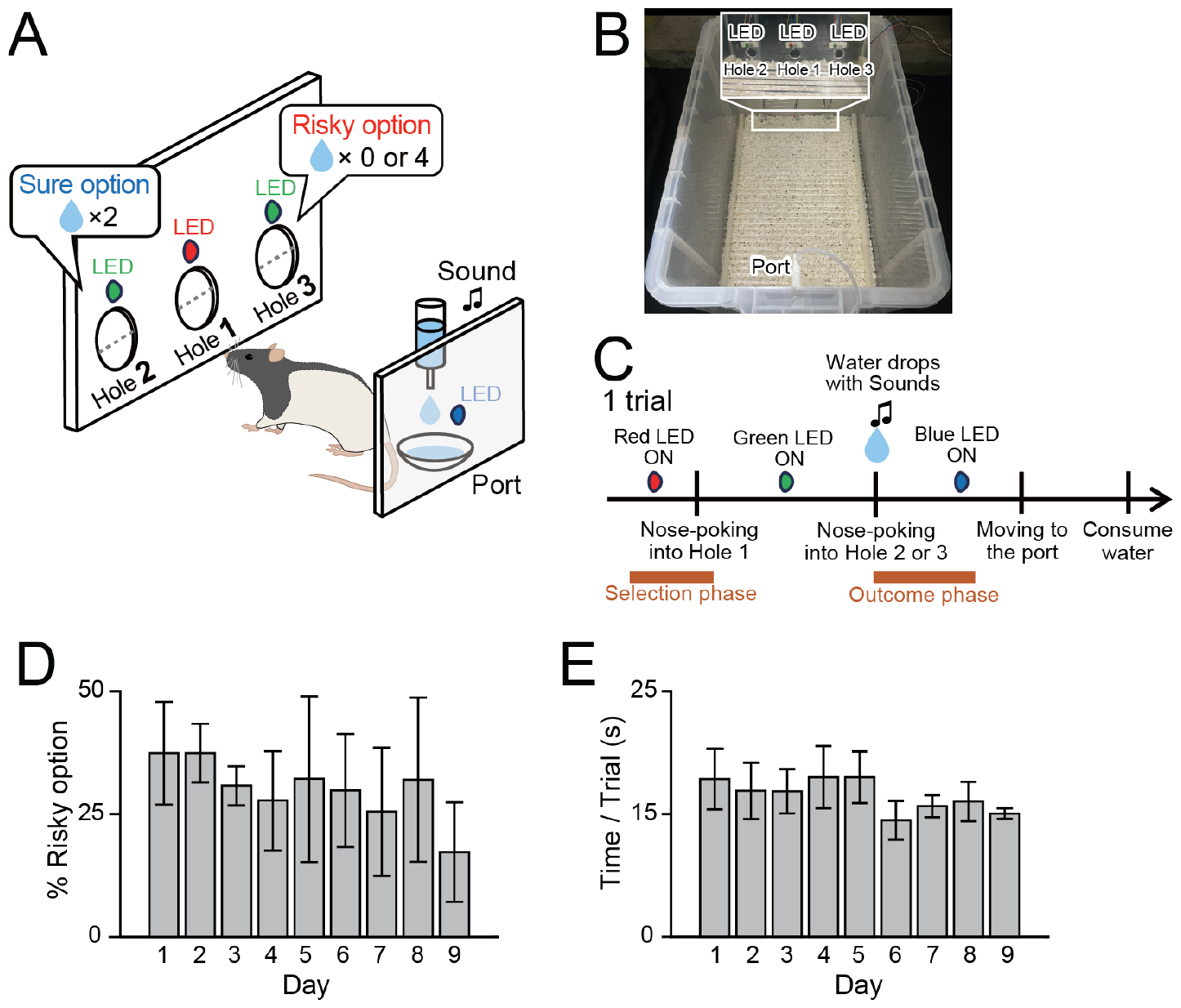
Performance of the gambling task. (A) Schematic illustration of the gambling task. (B) A representative picture of the apparatus. (C) A sequence of events in a trial of the gambling task. (D) The probability of selecting risky options in each day of the gambling task (*n* = 5 rats). Data are presented as the mean ± SEM. (E) The average duration spent for single trials in each day of the gambling task (*n* = 5 rats).

### Behavioral training

The amount of a water drop was 0.05 ml in all steps and the timing of the delivery of the water drops was indicated to the rats by clicking on a sound from the nozzle. The training consisted of the following steps. On the first day, each rat was placed in a box and trained to learn the position for water feeding. A water drop was automatically provided to the port every 10 s with a blue LED blinking for 1 s 100 times. The rats consumed all water drops. The next day, the rat was trained to receive a water drop on the port when the rat performed voluntary nose poking into hole 1 when the red LED above hole 1 blinked. The red LED was turned off when the rats performed valid nose poking. After water consumption, the rats had to leave the port for at least 2 s to initiate the next nose poking. The rats completed the task 200 times. The next day, the rat was trained to receive a water drop on the port when the rat first performed voluntary nose poking into hole 1 when the red LED above hole 1 was blinking and then voluntary nose poking into hole 2 when the red LED was turned off and the green LED above hole 2 was blinking. The green LED was turned off when the rat performed valid nose poking in hole 2. After the rat achieved this behavior 100 times, the rat performed the same procedure with the nose poked into hole 3 100 times. On the next two days, the rat was trained to receive a water drop on the port when the rat performed voluntary nose poking into hole 1 when the red LED above hole 1 was blinking, and then performed voluntary nose poking into holes 2 or 3 when the red LED was turned off and the green LEDs above both holes (holes 2 and 3) were simultaneously blinking, 200 times in total. The green LEDs were turned off when the rats performed nose poking in either hole 2 or 3. If the rat consistently chose nose poking on a specific side (hole 2 or 3) too frequently, training with the other side was extended for several days. From the 5th day, the rats were trained to perform the same behavior under the condition where one and two water drops were constantly provided on the port after they performed nose poking in holes 2 and 3, respectively. This training was repeated daily until the rat was able to choose hole 3 (containing a larger amount of water) in more than 70 out of 100 trials in three consecutive sessions. Achieving this performance criterion requires five to seven days. After meeting this criterion, five rats underwent surgery for electrode implantation as described later.

### Gambling task

After recovery from surgery, the rats underwent behavioral training for 1–2 d. After confirming that the rat had reached the performance criterion, as in the preoperative period, the rat started a gambling task. The task condition was similar to those in the training condition, except that choosing nose poking in the hole 2 constantly provided two water drops on the port, serving as “a sure option,” whereas choosing nose poking in hole 3 provided either zero or four water drops with a 50% probability each on the port, serving as “a risky option.” A trial was defined as the period from when the rat performed nose poking into hole 1 to when the rat returned to hole 1 after confirming the port. Irrespective of the number of water drops, the next trial was initiated. Each task session consisted of 100 trials. This task was repeated daily for up to 9 days.

### Surgery

Standard surgical procedures were similar to those described previously (Okonogi *et al*., 2018; Konno *et al*., 2019; Kuga *et al*., 2019; Konno *et al*., 2022). The rats were anesthetized with 1–2% isoflurane gas in air. Two ECG electrodes (stainless steel wires; AS633, Cooner Wire Company) were sutured to the tissue beneath the skin of the upper chest. The rats were then fixed in a stereotaxic instrument with two ear bars and a nose clamp. Two electromyography (EMG) electrodes were implanted in the dorsal neck area. All the wires were secured to the skull using dental cement. After completing all surgical procedures, anesthesia was terminated, and the rats were allowed to awaken from anesthesia. Following the surgery, each rat was housed with free access to water, food, and daily observations.

### Electrophysiological recording

Each rat was connected to the recording equipment via Cereplex M (Blackrock), a digitally programmable amplifier placed close to the head of the animal. The output of the headstage was obtained using a Cereplex Direct recording system (Blackrock Microsystems), a data acquisition system, via a lightweight multiwire tether and commutator. To record the electrophysiological signals, the electrical interface board was connected to a Cereplex M digital headstage, and the digitized signals were transferred to a Cereplex Direct data acquisition system (Blackrock Microsystems). Electrical signals were sampled at 2 kHz and low-pass-filtered at 500 Hz.

### Analysis

All analyses were performed using MATLAB (MathWorks). ECG traces were bandpass-filtered at 20–200 Hz, and beat-to-beat intervals (R-R intervals) were calculated from the timestamps of the R-wave peaks. Instantaneous heart rates were computed based on R-R intervals.

All data are presented as the mean ± SEM. For each statistical test, the data normality was first determined using the *F* test. Two-sample data were compared using a paired *t*-test. Multiple group comparisons were performed by Tukey-Kramer test after one-way ANOVA. The null hypothesis was rejected at *P* < 0.05 level.

## Results

### Rat performance in a gambling task

A schematic illustration of the gambling task is presented in Fig. 1A. In the training phase, five male Long Evans rats were trained to perform a behavioral sequence: nose poking into hole 1, nose poking into holes 2 (left) or 3 (right) in the wall of a box, and consumption of a constant number of water drops at a water port on the opposite wall inside the box (Fig. 1B) (for more details, see the Methods). Under all conditions, nose poking into holes 2 or 3 instantly triggered a clicking sound indicating the delivery of water drops at the port, so that the rats were notified whether water drops were delivered immediately after nose poking into holes 2 or 3. After the rats learned this behavioral pattern, they underwent surgery for heart rate recordings. After recovery from the surgery, the rats were tested using a gambling task. Choosing nose poking into hole 2 constantly provided two water drops on the port (termed a sure option), whereas choosing nose poking into hole 3 provided either zero or four water drops with a 50% probability each on the port (termed a risky option) (Fig. 1A). Thus, the expected value in each trial remained consistent regardless of which option was selected. In one trial, the period before nose poking into hole 1 and immediately after poking the nose out of hole 1 was considered a selection period, during which the rats likely decided which options to select (Fig. 1C). The period immediately after nose poking into holes 2 or 3 was considered as the outcome period, as the rats were notified by the clicking sound whether they could receive water drops. The rats were tested 100 times a day. On the initial day (day 1) of the gambling task, the rats learned for the first time the differences between sure and risky options, with probabilistic outcomes or losses. This day was excluded from further heart rate analyses. On the following days (from day 2), the probability of choosing risky options remained nearly constant at an average of 29.4 ± 3.9% (Fig. 1D), meaning that the rats became familiar with this task condition, and learned to balance this probabilistic choice. The duration for each trial stabilized at an average of 17.0 ± 2.0 s (ranging from 10.0 s to 21.7 s) from day 2 (Fig. 1E), verifying that the rats optimized their behavioral patterns without unnecessary movements.

### Gradual decreases in baseline heart rates were observed during the gambling task in a day

We analyzed how heart rates changed across trials in rats engaged in the gambling task (Fig. 2A). As the rats repeated the trials each day, we noticed that baseline heart rates at the end of the trials were significantly lower than those at the beginning of the trials. Plotting of all averaged heart rates (0−1.7 s before nose poking into hole 1) in every trial confirmed that heart rates gradually decreased from ∼450 bpm to ∼350 bpm as the trials progressed (Fig. 2B). In contrast, the probability of selecting risky options was almost constant, with no significant differences throughout the trials on each day (Fig. 2C; *F*_4,20_ = 0.21, *P* = 0.93, one-way ANOVA). These results suggest that the decreasing trend in heart rates across trials did not crucially affect the probability of animal selection.

**Figure 2.**
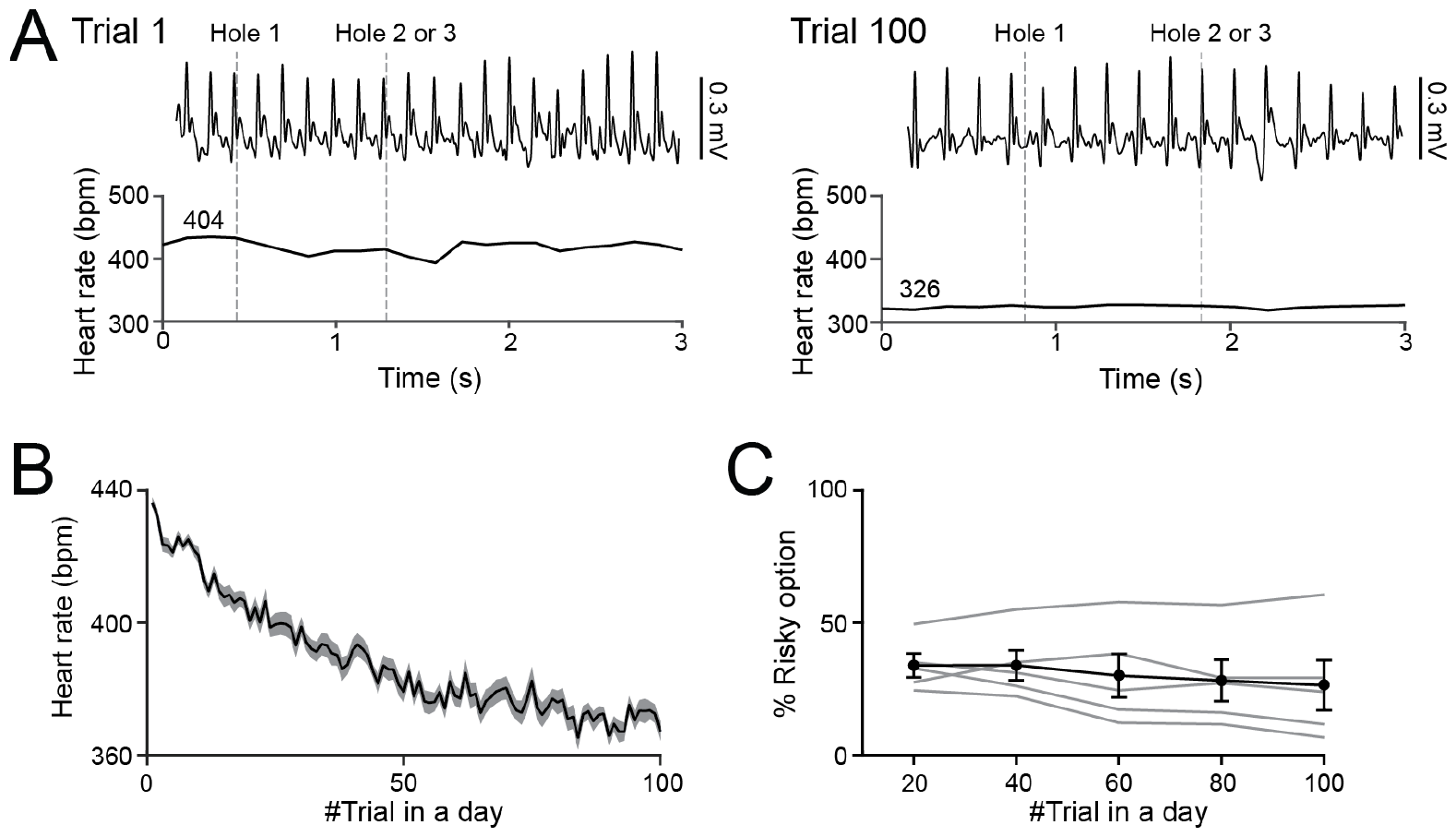
Changes in baseline heart rates across trials in a day. (A) Representative instantaneous heart rate changes in Trial 1 (left) and Trial 100 (right). Gray vertical lines represent the timing of nose poking into hole 1 and hole 2 or 3. Baseline heart rates (in bpm) are shown above the traces. (B) Changes in baseline heart rates over all 100 trials in each day of the gambling task. The shaded area represents the SEM. (C) Changes in the probability of selecting risky options in each day of the gambling task (*n* = 5 rats; bin = 20 trials). Data are presented as mean ± SEM.

### Dynamic heart rate changes in response to risky selections and outcomes

We analyzed how heart rates changed dynamically during the selection of certain or risky options in the task. Plotting instantaneous heart rates aligned with the onset of nose poking into hole 1, corresponding to the selection phase, revealed that heart rates significantly decreased after nose poking into hole 1, regardless of whether the rats subsequently chose certain or risky options (Fig. 3A; sure: *t*_4_ = 4.79, *p* = 0.0087; risky: *t*_4_ = 5.60, *P* = 0.0050, paired *t* test). Notably, the reduction in heart rate was significantly greater when the rats subsequently took the risky options than sure options (Fig. 3B; *t*_4_ = 5.70, *P* = 0.0047, paired *t* test). This result demonstrates that the rats were more likely to exhibit risky behavior when their heart rates decreased, suggesting a preference for risky decision-making during heart rate reduction.

**Figure 3.**
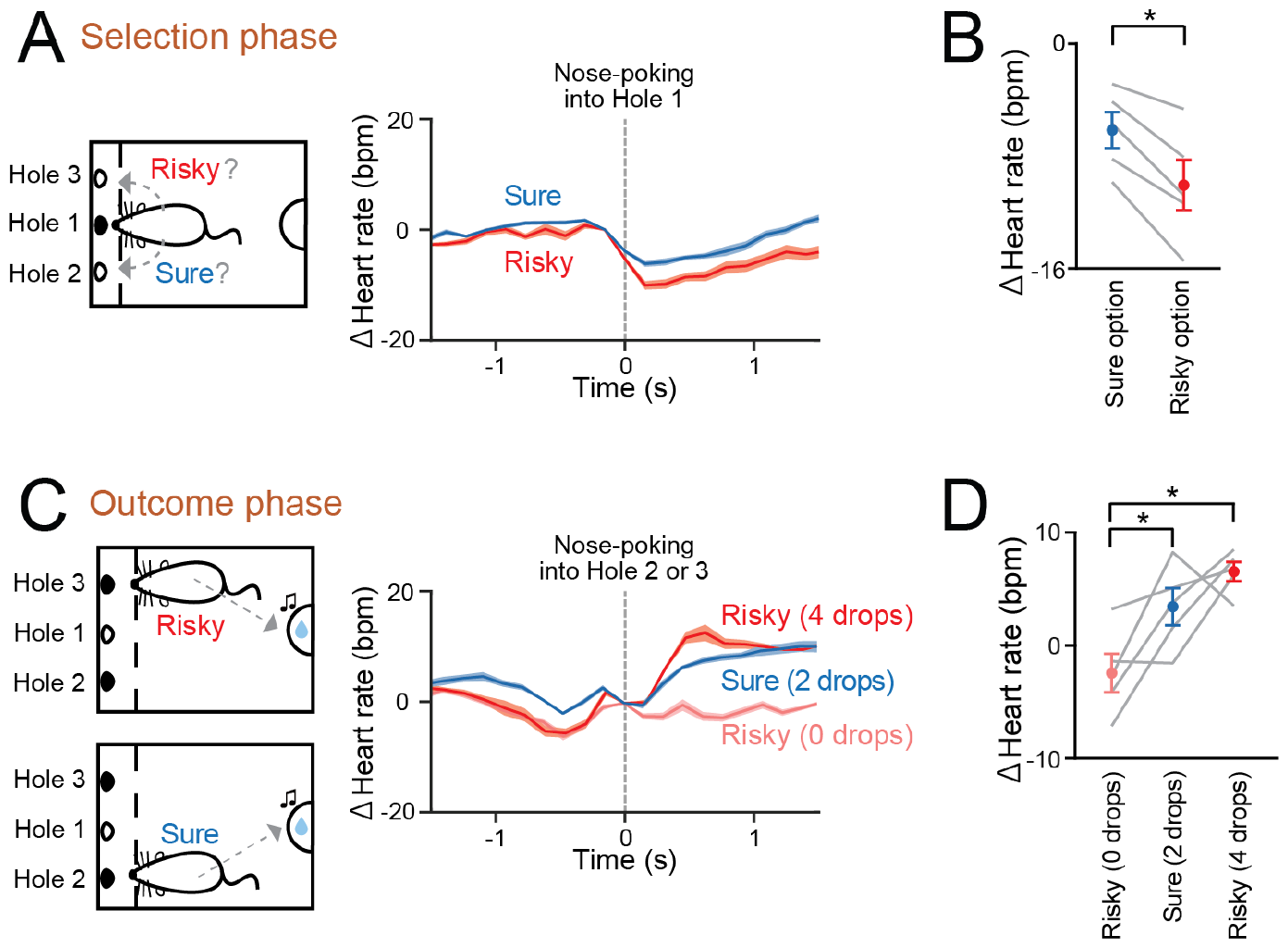
Heart rate changes during the selection phase and the outcome phase. (A) Changes in averaged heart rates aligned to the onset of nose poking into hole 1 (around the selection phase), separately plotted for the selection of sure (blue) and risky (red) options. The shaded area represents SEM. (B) Comparison of changes in averaged heart rates from –0.15 s to +0.15 s between the sure and risky options. Each gray line represents each rat. \**P* < 0.05, paired *t* test. (C) Changes in averaged heart rates aligned to the onset of nose poking into hole 2 or 3, corresponding with the timing of sounds signifying the amount of reward, separately plotted for when the amount of reward was 2 drops (blue) in the sure options, 4 drops (red), and 0 drops (thin red) in the risky options. The shaded area represents SEM. (D) Comparison of changes in averaged heart rates from 0 s to +0.3 s among when the rats that received 2 drops (blue) in the sure options, 4 drops (red) and 0 drops (thin red) in the risky options. Each gray line represents each rat. \**P* < 0.05, Tukey-Kramer test after one-way ANOVA.

Next, we analyzed how heart rates changed dynamically when the outcome was disclosed to the rats. Plots of the instantaneous heart rates aligned to the onset of nose poking into holes 2 or 3, corresponding to the outcome phase in the presence of clicking sounds (Fig. 3C). When the water amount was four drops in the risky options, significant increases in heart rates occurred during the outcome phase (*t*_4_ = −7.72, *P* = 0.0015, paired *t* test). On the other hand, such significant changes were not observed when the water amount was two drops in the sure options (*t*_4_ = −2.08, *P* = 0.11, paired *t* test) and no (zero) drops in the risky options (*t*_4_ = 1.44, *P* = 0.22, paired *t* test). Overall, heart rate increases were significantly larger with four drops in the risky options and with two drops in the sure options than with no drops in the risky options (*P* < 0.05, Tukey-Kramer test after one-way ANOVA) (Fig. 3D). These results demonstrate that the rats exhibited greater increases in their heart rates, depending on the outcomes of their risk-taking behaviors.

## DISCUSSION

In this study, we monitored gambling-related heart rate changes as rats performed a gambling task. On the day of recording, the baseline heart rates gradually decreased as the trials progressed, possibly representing acclimatization to the task condition. In the gambling task, instantaneous heart rates before selecting risky options were lower than those before selecting certain options. In addition, instantaneous heart rates after being notified of the outcome increased more as the number of outcomes increased.

Human studies have shown that healthy subjects display increased heart rates following wins, vs decreased heart rates after losses (Goudriaan *et al*., 2006). Our observations of outcome-related heart rate changes are consistent with those of this human study. In contrast, problem gamblers have decreased heart rates following both wins and losses, indicating a reduced responsiveness to gambling-related rewards (Goudriaan *et al*., 2006; Lole *et al*., 2014). Taken together, the rats tested in this study were considered to have become accustomed to typical gambling behavior and were not a model representing problem gamblers. In human studies, the manner in which heart rate is altered during gambling depends on different gambling frequencies or addiction levels (Leary & Dickerson, 1985; Coulombe *et al*., 1992; Griffiths, 1993; Goudriaan *et al*., 2006; Lole *et al*., 2014). In addition, few studies have examined how heart rates change dynamically across various periods of gambling, such as the anticipation of rewards and reception of outcomes. Interestingly, heart rate in humans has been observed to decrease rather than increase when anticipating future rewards during gambling (Goudriaan *et al*., 2006). Our observation of decreased heart rate in rats during the selection phase is consistent with this human study. Anticipation-related heart rate changes suggest that rats can effectively adapt to gambling environments by controlling their arousal levels.

Several possible physiological mechanisms are thought to underlie the tendency of heart rate reductions toward risky selection. The first explanation is that changes in heart rate simply reflect a byproduct of brain activity via efferent brain-heart transmission. In this scenario, ongoing activity in the brain first shifts to a state that preferentially triggers behavior toward gambling while, in parallel, attenuating autonomic sympathetic signals to reduce overall arousal levels and cardiac activity. It remains unknown whether there is a shared brain state that concurrently induces both risky selection and a reduction in arousal levels. The second explanation is that gambling-related behavior is influenced by heart-to-brain communication. The somatic marker hypothesis suggests that states of unconscious bodily arousal guide decision-making processes (Damasio, 1996). According to this hypothesis, heart rate reduction may facilitate gambling behavior by transferring afferent (interoceptive) signals from the heart to the brain (Craig, 2002). Consistent with this idea, recent studies have shown that acceleration of the heart rate by artificial cardiac stimulation increases anxiety-like behavior (Hsueh *et al*., 2023) and that heart rate changes are related to fearful behavior (Klein *et al*., 2021). Taken together, this suggests that decreased heart rate may effectively alleviate anxiety levels, thereby potentially favoring risky selection.

In conclusion, this study revealed dynamic changes in instantaneous heart rate in response to risky selections and outcomes during gambling. These insights are expected to aid in identifying expertise or acclimatization to gambling through physiological signals. Further studies are required to understand how these normal physiological signals are altered in animal models of decision-making deficits during gambling, such as addiction or psychiatric disorders.

## ACNOWLEDGEMENTS

This work was supported by KAKENHI (20H03545 and 21H05243) from the Japan Society for the Promotion of Science (JSPS), grants (JPMJCR21P1 and JPMJMS2292) from the Japan Science and Technology Agency (JST) to T. Sasaki, KAKENHI (23K11321) to A. Tamura, and a JSPS Research Fellowship to N. Kuga.

## AUTHOR CONTRIBUTIONS

F.F. A.T. and T.S. designed the study. F.F. and N.K. performed surgeries. F.F. acquired the behavioral and electrophysiological data and performed the analyses. F.F., A.T., and T.S. prepared the figures; T.S. wrote the main manuscript text. All the authors reviewed the main manuscript text.

## REFERENCES

Anderson, G. & Brown, R.I. (1984) Real and laboratory gambling, sensation-seeking and arousal. Br J Psychol, 75 (Pt 3), 401–410.

Barrus, M.M. & Winstanley, C.A. (2016) Dopamine D3 Receptors Modulate the Ability of Win-Paired Cues to Increase Risky Choice in a Rat Gambling Task. J Neurosci, 36, 785–794.

Coulombe, A., Ladouceur, R., Desharnais, R. & Jobin, J. (1992) Erroneous perceptions and arousal among regular and occasional video poker players. J Gambl Stud, 8, 235–244.

Coventry, K.R. & Hudson, J. (2001) Gender differences, physiological arousal and the role of winning in fruit machine gamblers. Addiction, 96, 871–879.

Coventry, K.R. & Norman, A.C. (1997) Arousal, sensation seeking and frequency of gambling in off-course horse racing bettors. Br J Psychol, 88 ( Pt 4), 671–681.

Craig, A.D. (2002) How do you feel? Interoception: the sense of the physiological condition of the body. Nat Rev Neurosci, 3, 655–666.

Damasio, A.R. (1996) The somatic marker hypothesis and the possible functions of the prefrontal cortex. Philos Trans R Soc Lond B Biol Sci, 351, 1413–1420.

Goudriaan, A.E., Oosterlaan, J., de Beurs, E. & van den Brink, W. (2006) Psychophysiological determinants and concomitants of deficient decision making in pathological gamblers. Drug Alcohol Depend, 84, 231–239.

Griffiths, M. (1993) Tolerance in gambling: an objective measure using the psychophysiological analysis of male fruit machine gamblers. Addict Behav, 18, 365–372.

Hsueh, B., Chen, R., Jo, Y., Tang, D., Raffiee, M., Kim, Y.S., Inoue, M., Randles, S., Ramakrishnan, C., Patel, S., Kim, D.K., Liu, T.X., Kim, S.H., Tan, L., Mortazavi, L., Cordero, A., Shi, J., Zhao, M., Ho, T.T., Crow, A., Yoo, A.W., Raja, C., Evans, K., Bernstein, D., Zeineh, M., Goubran, M. & Deisseroth, K. (2023) Cardiogenic control of affective behavioural state. Nature, 615, 292–299.

Ishii, H., Ohara, S., Tobler, P.N., Tsutsui, K. & Iijima, T. (2012) Inactivating anterior insular cortex reduces risk taking. J Neurosci, 32, 16031–16039.

Klein, A.S., Dolensek, N., Weiand, C. & Gogolla, N. (2021) Fear balance is maintained by bodily feedback to the insular cortex in mice. Science, 374, 1010–1015.

Konno, D., Ikegaya, Y. & Sasaki, T. (2022) Weak representation of awake/sleep states by local field potentials in aged mice. Sci Rep, 12, 7766.

Konno, D., Nakayama, R., Tsunoda, M., Funatsu, T., Ikegaya, Y. & Sasaki, T. (2019) Collection of biochemical samples with brain-wide electrophysiological recordings from a freely moving rodent. J Pharmacol Sci, 139, 346–351.

Kuga, N., Nakayama, R., Shikano, Y., Nishimura, Y., Okonogi, T., Ikegaya, Y. & Sasaki, T. (2019) Sniffing behaviour-related changes in cardiac and cortical activity in rats. J Physiol, 597, 5295–5306.

Leary, K. & Dickerson, M. (1985) Levels of arousal in high- and low-frequency gamblers. Behav Res Ther, 23, 635–640.

Lole, L., Gonsalvez, C.J., Barry, R.J. & Blaszczynski, A. (2014) Problem gamblers are hyposensitive to wins: an analysis of skin conductance responses during actual gambling on electronic gaming machines. Psychophysiology, 51, 556–564.

Lole, L., Gonsalvez, C.J., Blaszczynski, A. & Clarke, A.R. (2012) Electrodermal activity reliably captures physiological differences between wins and losses during gambling on electronic machines. Psychophysiology, 49, 154–163.

Okonogi, T., Nakayama, R., Sasaki, T. & Ikegaya, Y. (2018) Characterization of Peripheral Activity States and Cortical Local Field Potentials of Mice in an Elevated Plus Maze Test. Front Behav Neurosci, 12, 62.

Orsini, C.A., Moorman, D.E., Young, J.W., Setlow, B. & Floresco, S.B. (2015) Neural mechanisms regulating different forms of risk-related decision-making: Insights from animal models. Neurosci Biobehav Rev, 58, 147–167.

Tremblay, M., Silveira, M.M., Kaur, S., Hosking, J.G., Adams, W.K., Baunez, C. & Winstanley, C.A. (2017) Chronic D(2/3) agonist ropinirole treatment increases preference for uncertainty in rats regardless of baseline choice patterns. Eur J Neurosci, 45, 159–166.

